# The inactivation of DHHC7 in mouse liver promotes diet-induced obesity through a hepatic Prg4–GPR146 axis

**DOI:** 10.64898/2025.12.30.697005

**Authors:** Yingmin Sun, Ying Liu, Siyu Wang, Hanyu Wu, Xiaoli Hou, Jiaqi Duan, Junkai Pei, Yanhua Xu, Xiaoxiang Hu, Bing Chen

## Abstract

The liver secretes hepatokines that coordinate whole-body metabolism. Posttranslational lipidation of signaling proteins by DHHC palmitoyltransferases controls membrane targeting and signaling, yet the role of hepatic palmitoylation in systemic metabolic regulation is largely unexplored. Using an inducible, liver-specific DHHC7 knockout (D7LKO) mouse, we report that loss of DHHC7 in hepatocytes potentiates adenylyl cyclase–PKA–CREB signaling through reducing palmitoylation of inhibitory G protein α subunit (Gαi), leading to transcriptional elevation and secretion of proteoglycan 4 (Prg4). Elevated circulating Prg4 acts on adipocytes by binding GPR146 via Prg4’s N-terminal SMB domain, suppressing adipocyte PKA substrate phosphorylation and hormone-sensitive lipase (HSL) Ser563 phosphorylation, thereby impairing lipolysis. Under high-fat diet (HFD) feeding, D7LKO mice and mice with adenoviral hepatic Prg4 overexpression develop pronounced obesity characterized by increasing brown, subcutaneous, and visceral fat mass and adipocyte hypertrophy; strikingly, these animals show little impairment in glucose tolerance or circulating triglycerides but display elevated plasma cholesterol. Conditioned medium containing Prg4 recapitulates suppression of adipocyte HSL phosphorylation in vitro, and SMB-deleted Prg4 fails to bind GPR146 or inhibit HSL phosphorylation. Our findings define a liver palmitoylation– hepatokine axis that controls adipose lipolysis and predisposes to diet-induced fat accumulation, establishing Prg4–GPR146 as a mechanistic link between hepatic signaling and adipose energy mobilization.

## Introduction

Inter-organ communication via secreted factors is fundamental to systemic metabolic homeostasis. The liver is central to this network: it controls systemic glucose and lipid fluxes and secretes hepatokines that modulate peripheral tissues (1-3). Altered hepatokine signaling contributes to metabolic disease, but the regulatory mechanisms that govern hepatokine expression and secretion in response to intracellular signaling states are incompletely defined.

Protein S-palmitoylation catalyzed by DHHC family palmitoyltransferases targets proteins to membranes and modulates signaling complexes (4, 5). There are total of 23 members in DHHC protein family (6, 7). All DHHC proteins contain multiple transmembrane domains (from 4-6) and a DHHC motif. The transmembrane domains target the DHHC protein to cellular membranes and form a binding pocket for palmitoyl-CoA and DHHC motif constitutes the core catalytic site. Since DHHC palmitoyltransferase is a membrane associated enzyme, the palmitoylation reaction occurs either on or near the membrane. At present, the *in vivo* function of DHHC proteins largely remains unknown. However, because of the importance of palmitoylation in a wide array of biological processes such as cell signaling, metabolic reactions and vesicle trafficking (8), DHHC proteins are believed to play an important role in the regulation of cell function (9).

DHHC7 is a member of DHHC protein family. Current studies revealed that DHHC7 palmitoylates multiple substrates implicated in immune signaling (10, 11), trafficking (12), and metabolism (13, 14), yet its liver-specific role in whole-body energy balance has not been directly tested. Here we generated inducible, liver-specific DHHC7 knockout mice to probe how altered palmitoylation in hepatocytes impacts hepatokine output and systemic metabolism. We identify proteoglycan 4 (Prg4), a secreted matrix protein previously implicated in joint lubrication (15, 16) and emerging metabolic roles (17), as a DHHC7-regulated hepatokine. Mechanistically, DHHC7 maintains Gαi palmitoylation and inhibitory control over adenylyl cyclase (18, 19); DHHC7 loss attenuates Gαi palmitoylation, augments hepatic cAMP–PKA–CREB signaling, and elevates Prg4 expression. Circulating Prg4 binds adipocyte GPR146 to suppress PKA-dependent HSL phosphorylation and lipolysis, promoting adipose lipid accumulation under high-fat diet. These data reveal a palmitoylation-dependent hepatic control of a hepatokine that tunes adipocyte lipolysis and systemic fat accrual.

## Results

### Liver-specific DHHC7 deletion produces HFD-dependent obesity with increased adipose mass

To investigate the role of DHHC7 in the liver, we generated DHHC7 floxed mice (supplementary Figure 1-Fig. S1) and crossed them with Albumin-CreERT2 to produce inducible, liver-specific DHHC7 knockout animals (D7LKO) (Fig. S2). Then, tamoxifen was administrated to adult D7LKO mice as well as DHHC7 floxed mice (served as control:CON) for three times every other day. A week later, total liver RNA and extracts were prepared for quantitative RT-PCR and Western blot to determine the effectiveness of DHHC7 gene deletion. Shown in Fig. 1A and B, tamoxifen administration to D7LKO mice reduced hepatic DHHC7 mRNA and protein >90% relative to controls, an indication that DHHC7 is successfully deleted in D7LKO mice upon Tamoxifen administration.

**Figure 1.**
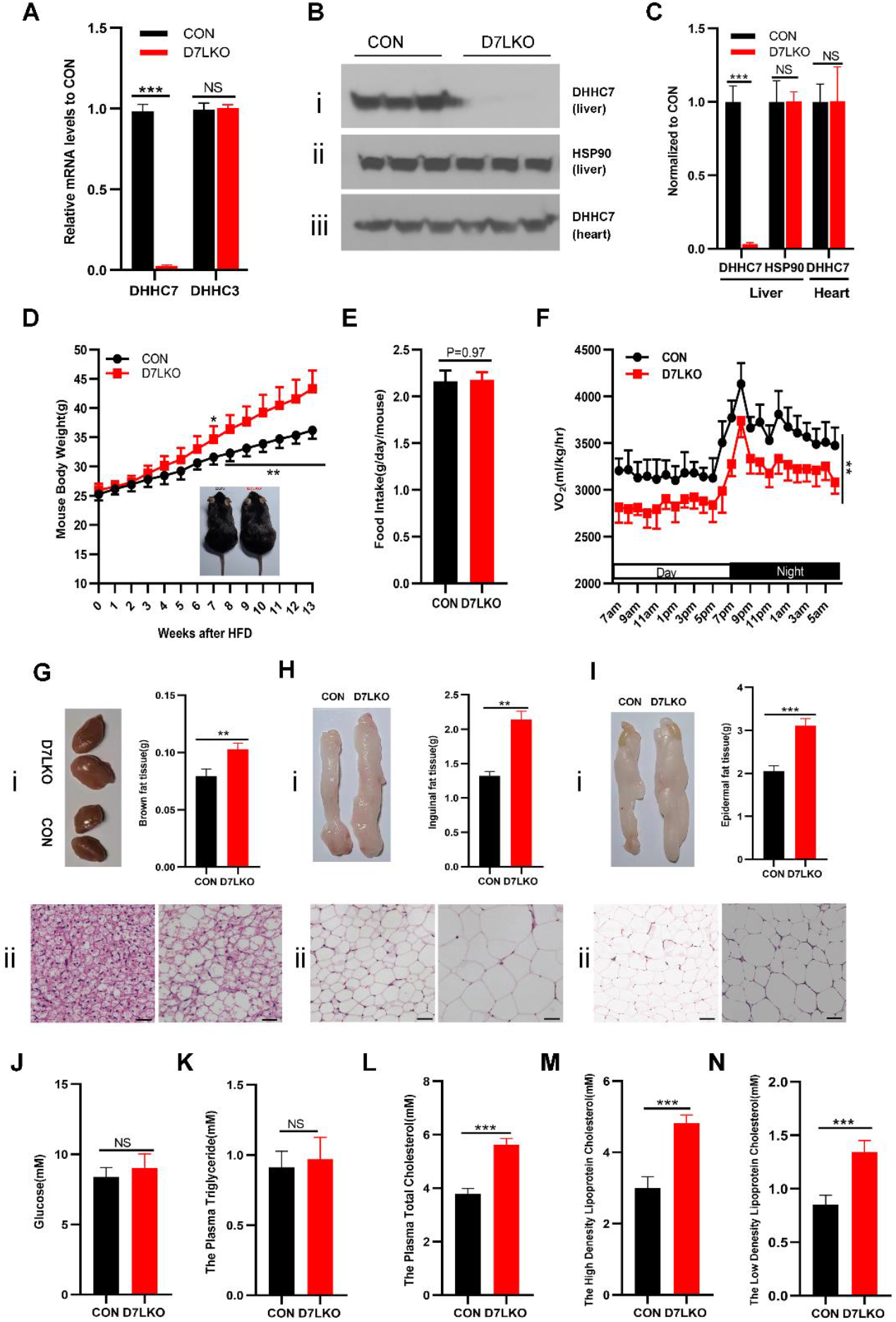
DHHC7 knockout in liver promotes obesity development and increases the plasma cholesterol under high fat diet feeding. A. Real-time PCR analysis the expression of DHHC3 and DHHC7 in liver from CON and D7LKO mice to show that DHHC7 expression is diminished in D7LKO mouse liver. B. Western blot of total liver extracts from D7LKO and CON mice with anti-DHHC7 (i), anti-Hsp90 (ii), respectively. Panel iii shows the expression DHHC7 in mouse heart. C Densitometric analysis of protein expression in B, the level of indicated protein was set to 1 in control. D. The growth curve of D7LKO mice and control mice. The mice (group of 6) were feed with High fat diet as indicated time and body weight were measured each week. E. Food intake of Control and D7LKO mice. F. The oxygen consumption of Control and D7LKO mice. G, H and I. The fat mass of Brown, Inguinal and epididymal fat of control and D7LKO mice (panel i); The right of panel i represents quantitative analysis of left panel i; The panel ii showed the H&E staining of each individual fat to show the increased adipocyte size in D7LKO mice. Scal bar is 25μM. J and K. The plasma glucose and triglycerides of Control and D7LKO mice. L, M and N. The plasma total cholesterol, high density lipoprotein and low density lipoprotein cholesterol of Control and D7LKO mice. In all plotting panels, the bar graphs represent Means+ S.D., n=3-6. ** p<0.01. ***p<0.001.

Once knowing that DHHC7 was successfully deleted in D7LKO mice, we examined the metabolic parameters of D7LKO mice. D7LKO mice showed no apparent alteration in blood glucose, lipid as well as cholesterol on standard diet (Fig. S 3 and 4), implying that DHHC7 expression in the liver has limited impact on body metabolism.

Since there were no apparent metabolic phenotype in D7LKO mice, we challenged male D7LKO mice with high-fat diet (HFD, 60% kcal fat), a common cause of obesity. Shown Fig. 1D, D7LKO mice exhibited accelerated weight gain beginning at 6 weeks and reaching about 20% greater body weight than controls by 13 weeks. The food intake was unchanged (Fig. 1E), whereas whole-body oxygen consumption was reduced (Fig. 1F) in D7LKO mice, indicating that D7LKO mice have a lowered energy expenditure.

To understand the nature of D7LKO obesity, we extracted all of adipose tissues including brown fat, inguinal fat and epididymal fat. First, we measured the fat mass. Shown in Fig. 1G, H and I, we observed a substantial increase in the mass of these fats (panel i). Next, we prepared sections of each fat and performed H&E staining. We observed marked increases in the size of each adipocyte (panel ii).

Insulin resistance is often associated with obesity in part due to increased circulating free fatty acids and chronic inflammation (20-22). With this in mind, we measured the blood level of fasting glucose, triglycerides and cholesterol. The fasting blood glucose and circulating triglycerides were similar between D7LKO and control mice (Fig. J and K), in spite of pronounced adiposity of D7LKO mice. However, a higher plasma level of total cholesterol including as well as high density lipoprotein and low density lipoprotein cholesterol was observed in D7LKO mice(Fig. L,M and N). Taken together, these data indicate that D7LKO were predisposed to diet induced obesity which has limited impact on insulin sensitivity (also see ITT and GTT in Fig. S5), but with potentially hypercholesterolemia. It is noted that these observations are not limited male mice. A similar phenotype was also observed with female D7LKO mice (Fig. S6).

### Hepatic DHHC7 loss elevates Prg4 expression and hepatic Prg4 overexpression phenocopies the obese phenotype

In the preceding studies, we observed that inactivation of DHHC7 in the liver predisposed the mice to diet induced obesity. We reasoned that this is likely due to that DHHC7 hepatic knockout altered the production of hepatokine (s) which act on the adipose tissue. To this end, we performed proteomic analysis of liver tissue from HFD-fed D7LKO and control mice. Shown in Fig. 2A, there were multiple proteins differentially regulated. Among them, Prg4 is marked because it is only secreted protein that was notably elevated in D7LKO plasma proteomic analysis (Fig. S7 A and B). To verify that Prg4 expression is indeed elevated in D7LKO liver, we carried out Western blot with total liver extracts from D7LKO liver. Shown in Fig. 2B. the hepatic Prg4 protein in D7LKO mice was ∼3-fold higher than controls. Next, we also examined the plasma level of Prg4 by ELISA. Shown in Fig. S7 C, the plasma level of Prg4 in D7LKO mice increased more than 2 folds. Taken together, these data demonstrated that inactivation of DHHC7 in the liver elevated the level of Prg4 protein.

**Figure 2.**
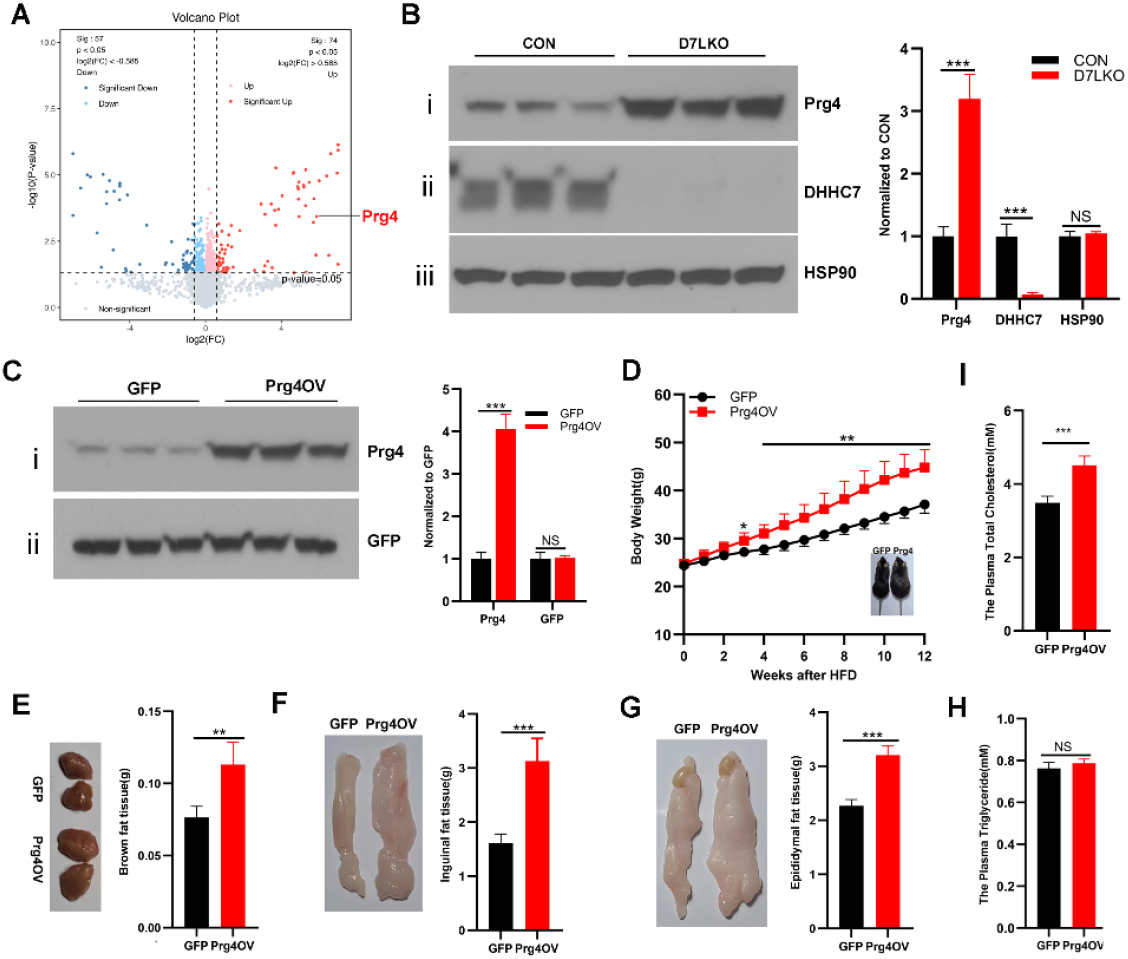
Inactivation of DHHC7 in the liver promotes the Prg4 expression and Overexpression DHHC7 in liver promotes the development of obesity under HFD feeeding. A. The Volcan plot of liver tissue proteomics of HFD male mice. The position of Prg4 is indicated by an arrow. B. Western blot of total liver extracts from Control and D7LKO mouse liver with anti-Prg4 (i), DHHC7 (ii) and HSP90(iii), respectively. The Right: Densitometric analysis of protein expression in the left, the level of indicated protein was set to 1 in control. C. Western blot of total liver extracts from the mice, which were administrated with empty and Prg4 adenoviral recombinant vector, with anti-Prg4 and anti-GFP antibodies, respectively. Right: Densitometric analysis of Prg4 expression in left, the level of indicated protein was set to 1 in GFP. The levels of DHHC7 were normalized to that of GFP encoding internally from adenoviral vectors. D. The growth curve of the mice (group of 6) administrated with either empty recombinant adenoviral vector or that encoding Prg4 under HFD feeding. The insert: the image of mice administrated with either empty recombinant adenoviral vector (GFP) or that encoding Prg4 (Prg4) under HFD feeding. E. F and G. The mass of Brown, inguinal and epididymal fat of the mice administrated with either empty recombinant adenoviral vector (GFP) or that encoding Prg4 (Prg4) after 12-week HFD feeding. H and I. The plasma level of triglycerides and total cholesterols in the mice administrated with either empty recombinant adenoviral vector (GFP) or that encoding Prg4 (Prg4) at 12 week HFD. Feeding. In all the plotting panels, the bar graphs represent Means+ S.D., n=3-6. ** p<0.01. ***p<0.001.

Because Prg4 level was elevated in D7LKO mice, we assumed that elevated Prg4 is a contribution factor for diet induced obesity of D7LKO mice. To test this hypothesis, we generated an adenoviral vector that expresses Prg4. Then, the emptied adenoviral vector or the one that encoding Prg4 were administrated to mice and the mice were challenged with high fat diet. Shown in Fig. 2C, adenoviral Prg4 administration increased hepatic Prg4 protein about 4 folds, comparable to levels comparable with D7LKO mice (Fig. 2B). Under HFD, adenoviral Prg4 administration drove marked body weight gain (∼27% more than controls) (Fig. 2D) and increased brown, inguinal, and epididymal fat mass (Fig. 2E, F and G) without altering circulating triglycerides but elevated total cholesterol (Fig. 2H and I), recapitulating key features of the D7LKO phenotype. Collectively, these data demonstrate that DHHC7 inactivation elevated Prg4 expression which constitutes a major factor to promote diet-induced adipose expansion.

### DHHC7 controls hepatic PKA–CREB activity through Gαi palmitoylation to regulate Prg4 transcription

Analyzing Prg4 expression revealed that CREB is a major transcriptional factor that modulates Prg4 expression (23, 24). CREB is a signal dependent transcription factor and its activation depends on its phosphorylation at Ser133 by PKA (25-27). Therefore, we assumed that the enhanced Prg4 expression by DHHC7 inactivation is through hepatic PKA–CREB pathway. To test this, the total liver extracts were prepared from D7LKO and control mice, respectively and subjected to Western blot with either phospho-PKA substrate antibodies or anti-phospho-CREB at Ser133 antibodies. Shown in Fig. 3A, D7LKO liver displayed markedly increased phosphorylation of PKA substrates (compare lane 1, 2, 3 with 4, 5, 6). Shown in Fig. 3B, DHHC7 inactivation in the liver led a three-fold increase in phospho-CREB at Ser133 (panel i) without altering total cellular CREB (panel ii), which is concurrent with increased Prg4 protein (panel iii).

**Figure 3.**
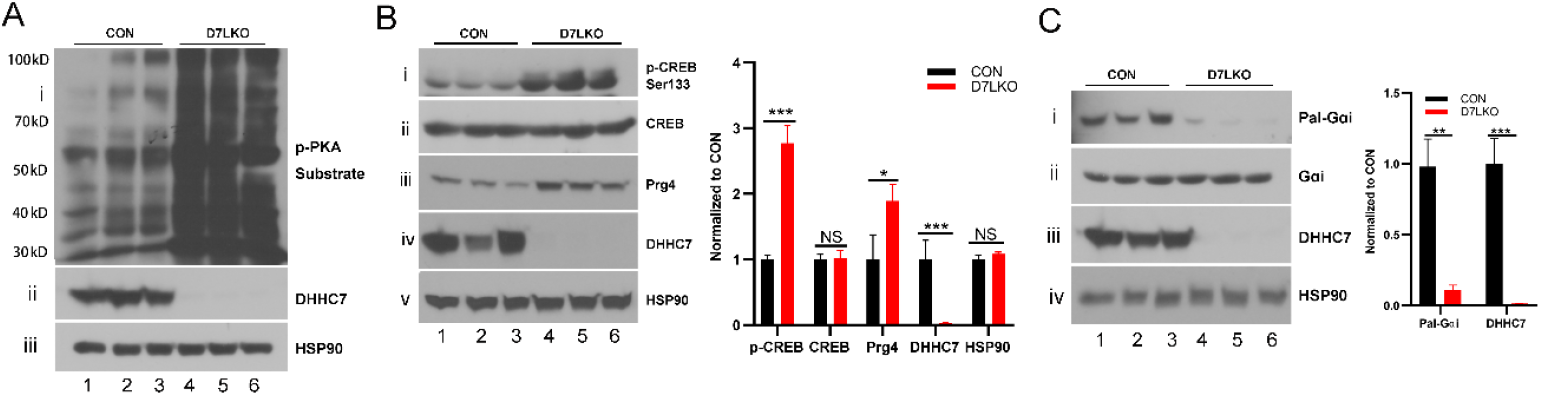
Inactivation of DHHC7 in the liver enhances PKA-CREB pathway. Western blot of total liver extract from either control or D7LKO mice with anti-phospho-PKA substrate (i), DHHC7 (ii) and HSP90 (iii) antibodies, respective. The Molecular Weight is marked at left in panel i. B. Western blot of total liver extract from either control or D7LKO mice with anti-phospho-CREB at Ser133 (i), CREB (ii), Prg4 (iii), DHHC7 (iv) and HSP90 (v) antibodies, respectively. The right: the densitometric analysis of the expression of each protein in right. The level of each protein in control was set as 1. C. TPC assay of Gαi in the liver protein from either control (con) or D7LKO. Panel i. palmitoylated. Panel ii, iii and iv show the level of total Gαi、 DHHC7 and HSP90 in total liver extracts. Left: Densitometric analysis of relative levels of palmitoylated Gai and DHHC7 in that the total level for each protein in control mouse liver was set as 1. In all the plotting panels, the bar graphs represent Means+ S.D., n=3. * p<0.05. ** p<0.01. ***p<0.001.

PKA is activated by cAMP, generated by adenylyl cyclase (AC) which is tightly regulated by G protein alpha in that Gαs stimulates whereas Gαi suppresses adenylyl cyclase activity (28). Gαi inhibitory activity on AC depends on its membrane location which is regulated by palmitoylation (18, 29-31). With this in mind, we assayed Gαi palmitoylation in livers using thiopropyl-Sepharose capture. Palmitoylated Gαi was substantially reduced in D7LKO livers while total Gαi levels were unchanged (Fig. 3C). A commonly used AC agonist is forskolin. We therefore examined whether Prg4 has any impact on forskolin induced CREB phosphorylation and found that pretreatment of Prg4 had no impact on forskolin induced CREB phosphorylation at Ser133 (Fig. S8), providing the evidence that DHHC7 acts on the upstream of AC. Since Gαi palmitoylation is required for its inhibitory activity, the loss of Gαi palmitoylation would impair inhibitory control of adenylyl cyclase, elevating cAMP → PKA activity and CREB-dependent Prg4 transcription.

### Circulating Prg4 suppresses adipocyte PKA signaling and HSL phosphorylation via GPR146 binding

In the preceding studies, we showed that inactivation of DHHC7 in the liver predisposed mice to diet induced obesity concurred with adipocyte hypertrophy. However, diet induced D7LKO mouse obesity is not associated with insulin resistance induced by free fatty acids released from adipose tissue (22). This implies that elevated Prg4 in D7LKO mice is likely involved in the adipocyte pathway that regulates lipolysis. With this in mind, we assessed adipose HSL phosphorylation at Ser563, an essential event for lipolytic activation (32-34), in the different adipose tissue from D7LKO mice. Shown in Fig. 4A, B and C. phospho-HSL (Ser563), was reduced >90% in brown, inguinal, and epididymal fat depots despite unchanged total HSL protein in D7LKO mice.

**Figure 4.**
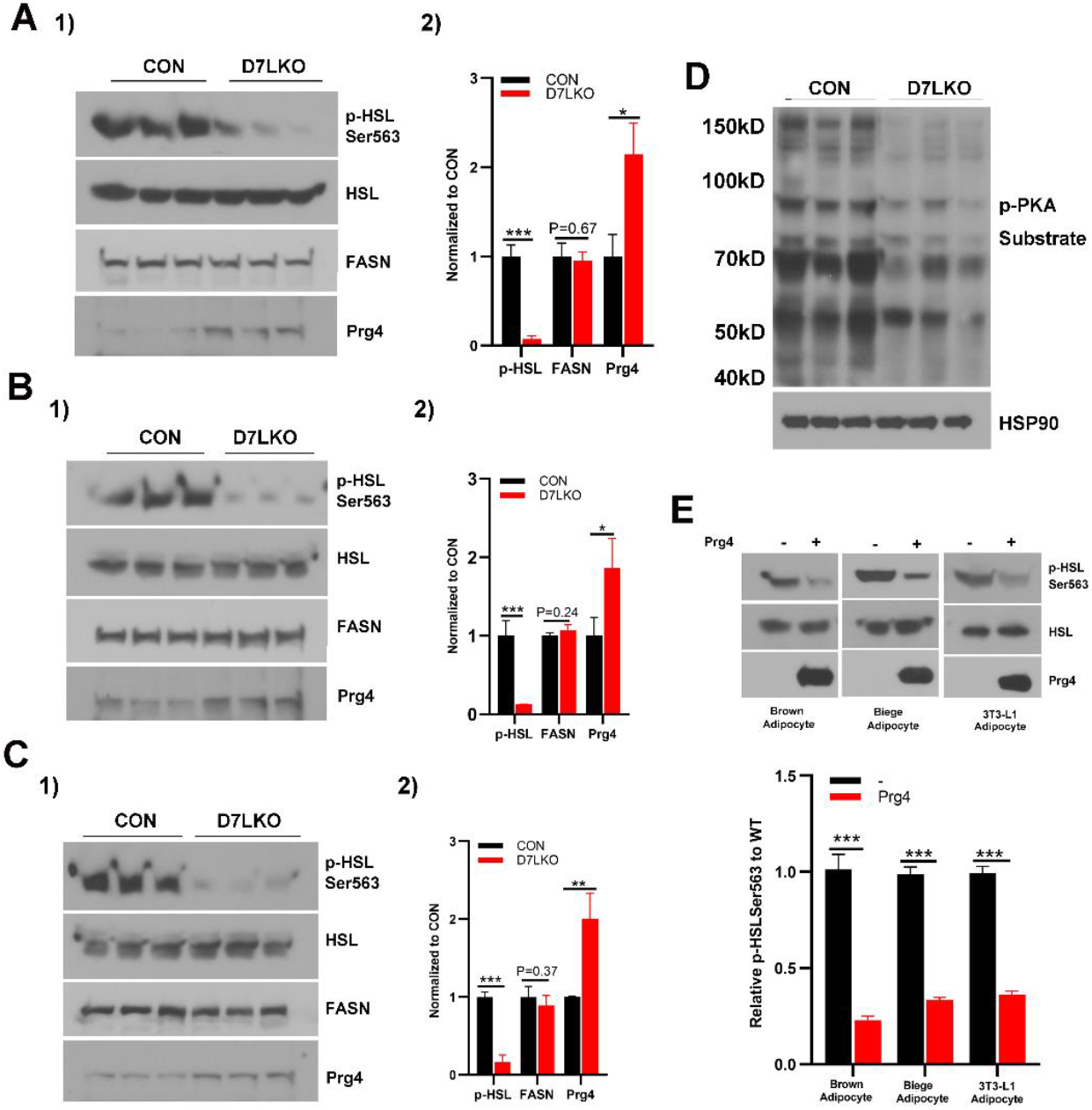
Prg4 suppresses adipocyte PKA signaling and HSL phosphorylation. A. 1) Western blot of total brown fat tissue lysate from control or D7LKO mice with anti-phospho-HSL at Ser563 (i), HSL (ii), Prg4 (iiii) and FASN antibodies, respectively. 2). Densitometric analysis of the level of indicated proteins in 1). The levels of each protein in control mice were set as 1. B and C. The same as A except Inguinal and Epididymal fat, respectively. D. Western blot of total brown fat extracts from control or D7LKO mice with anti-phospho-PKA substrate antibodies to show that inactivation of DHHC7 in the liver inhibited PKA substrate phosphorylation in adipocytes. E. The treatment of adipocytes with Prg4 suppresses HSL phosphorylation at 563. The differentiated Brown, Biege and 3T3-L1 adipocytes were incubated with the culture medium prepared from 293T cells expressing with or without Prg4 (see Material and Methods). 4 hours later, the total cell lysates were prepared for Western blot with anti-phospho-HSL at Ser563 (i). HSL (ii) and Prg4 (iii), respectively. 2) Densitometric analysis of indicated protein in 1). The level of indicated protein was set to 1 in control cells. In all plotting the panels, the bar graphs represent Means+ S.D., n=3. * p<0.05. ** p<0.01. ***p<0.001.

HSL phosphorylation at Ser563 is mediated by PKA in adipocyte (35). The view that the reduction of HSL phosphorylation at Ser563 implies that PKA signaling in D7LKO adipose tissue is impaired. To test this, we assessed the status of PKA substrates in D7LKO brown fat as a representative. Shown in Fig. 4D, adipose PKA substrate phosphorylation was similarly decreased. To direct test the impact of Prg4 on HSL phosphorylation at Ser563, conditioned medium from Prg4-expressing HEK293T cells was applied to differentiated brown, beige, and 3T3-L1 adipocytes for 8 h and the status of HSL phosphorylation at Ser563 was assessed in Western blot. Shown in Fig. E, Prg4 conditioned medium reduced HSL Ser563 phosphorylation more than 60% comparing to control medium, demonstrating that secreted Prg4 is sufficient to suppress adipocyte PKA–HSL signaling.

Prg4 is an extracellular protein (24). In adipocytes, PKA is activated by G protein-coupled receptors (GPCRs) (36). We reason that if anything, Prg4 likely acts on GPCR on the adipocyte surface to modulate PKA signaling. There are more than dozens of GPCRs in adipocytes, among them, GPR146 has been implicated in adipocyte lipolysis and hepatic cholesterol metabolism (37-40). With this in mind, we examined the interaction between Prg4 and GPR146 in co-immunoprecipitation assay with Flag-tagged GPR146 and Prg4. Specifically, Prg4 was expressed in 293T cells with or without co-expressing Flag-GPR146. Then, total cell lysates were prepared for immunoprecipitation with anti-Flag M2 affinity gel and anti-Flag M2 immunoprecipitates were analyzed in immunoblotting with anti-Prg4 antibodies. Shown in Fig. 5A, Prg4 was only detected in anti-Flag M2 immunoprecipitates at the present of Flag-GPR146 (compare lane 1 and 2, panel i). The differences were not because of Prg4 sample variation as comparable levels of Prg4 were seen in each sample.

**Figure 5.**
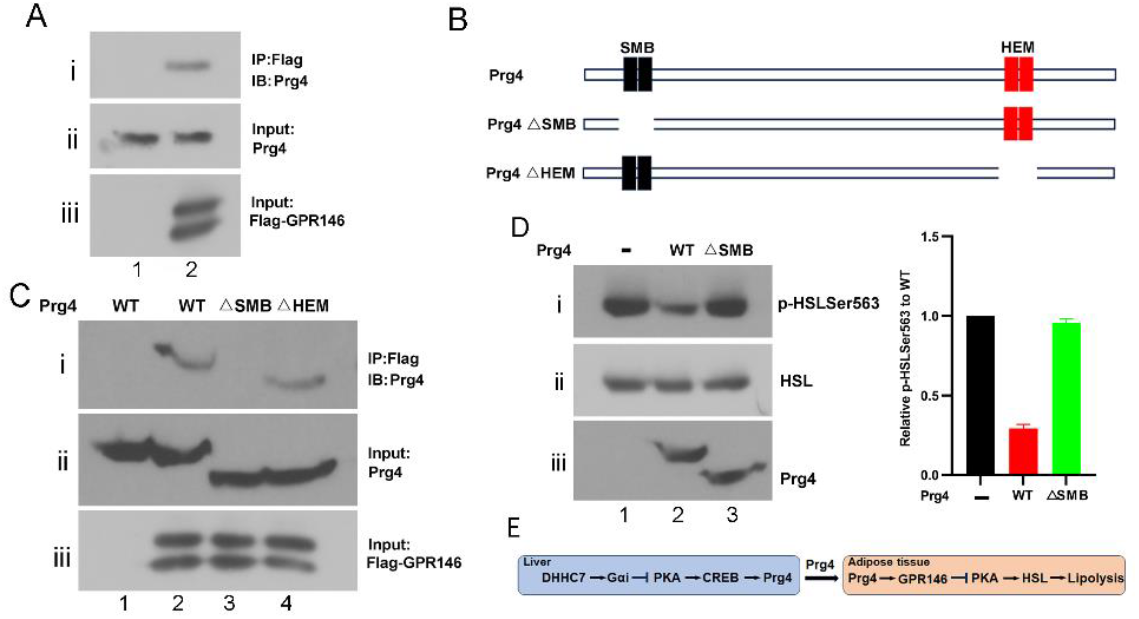
Prg4 interacts with GPR146 and this interaction is required for Prg4 to suppress HSL phosphorylation at S563. A. Co-immunoprecipitation assay of Flag tagged GPR146 and Prg4 co-expressed in HEK293T cells with anti-M2(Flag) affinity gel. Anti-M2 immunoprecipitates were probed with anti-Prg4 antibodies (i). Panel ii and iii showed the input levels of Prg4 and Flag-GPR146, respectively. B. Schematic representation of Prg4 structure and its two mutants. The two major domains of Prg4: SMB and HEM were indicated. C. Co-immunoprecipitation assay of the interaction between GPR146 with different forms of Prg4 to show that SMB domain of Prg4 mediates the interaction between Prg4 and GPR146. The experiments were performed essentially as A, except two Prg4 mutants were also used. ΔSMBPrg4: SMB domain was deleted. ΔHEMPrg4: HEM domain was deleted. D. The interaction between Prg4 and GPR146 is required for Prg4 to suppress HSL phosphorylation at Ser563. The experiments were carried out similar to Fig. 4E except ΔSMBPrg4 and ΔHEMPrg4 were also used. E The work model of Prg4 in liver and adipose tissue. In plotting panels, the bar graphs represent Means+ S.D., n=3. * p<0.05. ** p<0.010. ***p<0.001.

Prg4 consists of two functional domains including a N-terminal somatomedin-like domain (SMB) and a C-terminal hemopexin-like domain (HEM), which are proposed to mediate protein-protein interaction (24) (Fig. 5B). To understand the nature of the interaction between Prg4 and GPR146, we deleted either SMB or HEM domains in Prg4 and examined the role of each domain in the association of Prg4 with GPR146. Shown in Fig.5C of the co-immunoprecipitation assay of Flag-GPR146 and different forms of Prg4, Prg4 was easily detected in anti-Flag-GPR146 immunoprecipitates as well as that of ΔHEMPrg4 in which HEM domain was deleted (lane 2. and lane 4, panel i). In a sharp contrast, no Prg4 was detected in the immunoprecipitates of anti-Flag Δ SMBPrg4 in which SMB domain is deleted. Taken together, these data indicate that SMB domain of Prg4 mediates the interaction between Prg4 and GPR146.

We tested the potential association between Prg4 and GPR146 in co-immunoprecipitation assay. Co-immunoprecipitation studies in HEK293T cells revealed that Prg4 physically associates with Flag-tagged GPR146. Domain mapping showed that the N-terminal somatomedin-like (SMB) domain of Prg4 is necessary for binding: SMB-deleted Prg4 (DSMB) failed to co-precipitate with GPR146 whereas HEM-deleted Prg4 (DHEM) retained binding. Functional assays using conditioned media demonstrated that WT Prg4 and Δ HEM Prg4 suppressed adipocyte HSL Ser563 phosphorylation, but Δ SMB Prg4 did not(Fig. 5D). These results indicate that Prg4 reduces adipocyte PKA signaling and lipolysis through direct engagement of GPR146 via its SMB domain.

## Discussion

This study identifies a liver palmitoylation → hepatokine axis that regulates adipose lipolysis and predisposes to diet-induced obesity. DHHC7 activity in hepatocytes maintains Gαi palmitoylation, preserving inhibitory control on adenylyl cyclase. Loss of DHHC7 diminishes Gαi palmitoylation, elevates hepatic cAMP–PKA–CREB signaling, and promotes transcriptional upregulation and secretion of Prg4. Circulating Prg4 engages adipocyte GPR146 via its SMB domain, suppresses adipocyte PKA substrate phosphorylation and HSL Ser563 phosphorylation, reduces lipolytic flux, and thereby promotes fat accumulation under HFD. In all, DHHC7 knockout in liver enhances Prg4 expression, and Prg4 interacts with GPR146 in adipose tissue to inhibit lipolysis as described in Fig.5E.

Several aspects of the phenotype are notable. First, D7LKO-driven obesity occurs without overt insulin resistance or hypertriglyceridemia, distinguishing this form of adipose expansion from obesity caused by classical insulin resistance-driven lipotoxicity. Suppression of adipocyte lipolysis by Prg4 likely reduces systemic free fatty acid flux, explaining preserved glucose tolerance despite increased adiposity. Second, elevated plasma cholesterol in D7LKO and Ad-Prg4 mice suggests additional roles for Prg4– GPR146 signaling in hepatic cholesterol handling or intestinal lipid absorption; GPR146 has previously been implicated in cholesterol regulation, consistent with our findings. Third, the identification of Prg4 as a hepatokine expands its known biology beyond joint lubrication and suggests potential broader physiological and pathological roles.

Mechanistically, our data implicate Gαi palmitoylation as a critical control point linking hepatic palmitoylation state to transcriptional programs. While the palmitoylation capture assay demonstrates reduced Gαi palmitoylation in D7LKO liver, further biochemical work will be needed to define whether Gαi is a direct substrate of DHHC7 or whether other DHHC7 targets modulate Gαi palmitoylation or membrane targeting. Likewise, the downstream signaling dynamics between hepatic PKA–CREB activation and selective induction of Prg4 versus other CREB targets merit further dissection.

Initially identified as a joint lubricant, Prg4 has been implicated in tissue homeostasis regulation, inflammation as well as (41). In this report, we identified Prg4 as a hepatokine, providing a new insight into Prg4 function.

GPR146 has been implicated in the liver cholesterol metabolism and adipose lipolysis and proposed as a potential therapeutic target for modulating cholesterol homeostasis (42). Our studies here suggest that the Prg4–GPR146 axis provides a testable target: antagonizing Prg4 binding to GPR146, blocking GPR146, or restoring hepatic DHHC7 activity could potentially increase adipose lipolysis and limit diet-induced fat accumulation. Conversely, harnessing Prg4’s anti-lipolytic property might be useful where lowering circulating free fatty acids is desirable. Detailed evaluation of organ-specific roles and long-term metabolic consequences will be essential.

In summary, our work links hepatic palmitoylation to hepatokine output and adipose lipolysis, identifying DHHC7 and Prg4–GPR146 as key regulators of diet-induced adiposity. At present, the reason for hypercholesterolemia in D7LKO obese mice is not understood. Two possible scenarios: 1) DHHC7 modulates SREBP-2 in the liver and 2) Prg4 affects GPR146 in the liver, both of which are under investigation.

## Materials and Methods

### Animals

DHHC7 floxed mice (DHHC7^F/F^ mice) in which the exons 5 and 6 of the DHHC7 gene flanked by loxp sites using the CRISPR-Cas9 technique, were established by Shanghai Model Organisms Center, Inc. The genotyping primers are as following: 5’LoxpF-GTGTGCATTTGAAGTGTCTGCTG, 5’LoxpR - TGCCAACCCGAGAGCTTCCAAG; 3’LoxpF-GATCAGAATGTGGCAGCTTCATC, 3’LoxpR – CACAAGTTAGCCATGGACTGAGG. To generate liver specific inducible DHHC7 knockout mice, DHHC7 floxed mice were crossed with Albumin-CreERT2 (from Shanghai Model Organisms Center, Inc) to generate DHHC7 floxed Albumin-CreERT2 (DHHC7^F/F-AlbCREERT2^). To delete DHHC7 in liver, DHHC7^F/F-AlbCREERT2^ mice was oral gavage Tamoxifen (2g/kg, three times, one time for every other day) at six to eight weeks old. The mice were bred and housed at the Hebei University animal facility. Both male and female mice were used for experiments. All animal experiments were approved by the Institutional Animal Care and Use Committees (IACUC) of Hebei University and performed in accordance with all relevant guidelines and regulations of the IACUC of Hebei University. The mice were fed with normal chow diet (NCD) or high fat diet (HFD, 60% fat, Research Diets, Inc., D12492). The body weight were monitored weekly. The food intake was measured every day for individually caged mice for one week. Body composition was determined by nuclear magnetic resonance (Echo MRI). The O_2_ consumption, CO_2_ production and physical activity were measured in metabolic cages (Columbus Instruments). To examine the adipose tissue growth, different adipose tissues were removed from mice and weighted. Then, the adipose tissues were dissected and fixed in 10% neutral buffered formalin and sectioned for H&E Staining, which was performed in the facility core of Zhejiang Chinese Medical University. For glucose tolerance test (GTT), glucose was intraperitoneally injected at 1g/kg body weight after 12 hours fasting. For insulin tolerance test (ITT), human insulin were intraperitoneally injected at 0.75U/kg body weight after 6 hours fasting. Glucose level was determined with tail blood at indicated time after glucose or insulin injection using a One Touch ultra2 glucose meter as described (12). Serum triglycerides, total cholesterol, high density lipoprotein cholesterol, low density lipoprotein cholesterol and Prg4 were measured with triglyceride determination kit (A110-1-1, Nanjing Jiancheng Bioengineering Company), total cholesterol assay kit(A111-1-1, Nanjing Jiancheng Bioengineering Company), high-density lipoprotein cholesterol assay kit(A112-1-1, Nanjing Jiancheng Bioengineering Company), low-density lipoprotein cholesterol assay kit(A113-1-1, Nanjing Jiancheng Bioengineering Company),Prg4 ELISA kit (RK09131, ABclonal).

### Reagents and antibodies

All common chemicals, buffers were from Sigma, Thermo fisher scientific, Cayman biochemical and MCE. The antibodies used in the present study are as the following Anti-Flag (Cat#F3165) and anti-Flag M2-Affinity gel (Cat#A4596) from Sigma. Anti-HSP90 (Cat#4877), Anti-Phospho-HSLSer563 (Cat#4139), Anti-HSL (Cat#4107), Anti-Phospho-PKA Substrate (Cat#9624), Anti-Phospho-CREB Ser133 (Cat#9198), Anti-CREB(Cat#9197), from Cell Signaling Technology. Anti-FASN (Cat#ST1549) from Millipore. Anti-DHHC7 (Cat# A17981), Anti-Prg4(Cat#A15379), from Abcolonal.

### Plasmid and viral vector construction and production

The Flag-GPR146 was purchased from SinoBiological(Cat HG12959-CF). To generate lentiviral vector expressing Prg4, lentiviral vector pLV-GFP was digested with SalI, and blunted by T4 DNA polymerase, then digested by BamHI. Prg4 cDNA was amplified by PCR with primers: 5’-GAAGATCTATGGGGTGGAAAATACTTCC-3’ and 5’-TCAAGGACAGTTGTACCAGA TTTTG-3’, and the PCR product was digested with BglII. Then, the digested Prg4 cDNA was ligated with the above digested pLV-GFP.

The generation of mutant Prg4 in which either SMB or HEM was deleted were carried out with overlapping PCR with following corresponding primers: SMB domain deletion: Forward, 5’-GAAGATCTATGGGGTGGAAAATACTTCC and backward. 5’-TGGTGATGTG GGATTTTGAGATGAAACCTGTTG-3’, Forward:5’-AATCCCACATCACCATCTC-3’ and backward,5’-TCAAGGACAGTTGTACC AGATTTTG-3’; HEM domain deletion: Forward, 5’-GAAGATCTATGGGGTGGAAAATACTTCC-3’ and backward, 5’-GAGATGGTGATGTGGGATT-3’. Forward, 5’-AATCCCACATCACCATCTC-3’ and backward, 5’-TCAAGGACAGTTGTACCAGATTTTG-3’.To generate adenoviral vector encoding Prg4, its PCR product was digested by BglII, and inserted into BglII and EcoRV of pAdtrackTBG. The adenoviruses were produced and purified by a CsCl gradient as described (43).

### Cell cultures and Transient transfection

HEK293T were grown in high glucose Dulbecco’s modified Eagle’s medium (DMEM) supplemented with 10% (v/v) fetal bovine serum(FBS), 2mM L-glutamine, 100 units/ml penicillin and 100 μg/ml streptomycin (Invitrogen). The cells were transiently transfected with lipofectamine 2000(Invitrogen) according to manufacturer’s instruction.

To produce the culture medium containing different form of Prg4, empty lentiviral vector or lentiviral vectors encoding different forms of Prg4 were transiently transfected into 293T cells. 36 hours posttransfection, the cell culture medium was collected and filtered through 0.45μm syringe filter.

## Adipocyte Differentiation

For the 3T3-L1 preadipocyte differentiation (44), 3T3-L1 preadipocytes were cultured for two additional days after reaching 100% confluence and treated with differentiation medium (Dulbecco’s modified Eagle’s medium–high glucose containing 10% fetal bovine serum, 2.5 μg/ml insulin, 0.5 mM IBMX, 2.5 mM dexamethasone, 2 mM L-glutamine, 100 units/ml penicillin, and 100 μg/ml streptomycin) for 4 days. Then, the medium was changed to regular medium. After 7 days of differentiation, the adipocytes were used for experiments.

For the brown preadipocytes differentiation: the interscapular brown fat pad was dissected from the new born mice(day 1) and digested for 40 minutes with collagenase solution at 37°C. Then, digested tissue were filtered through 100 μm filters and spin at 1000rpm for 10 minutes. The pellet is resuspended and cultured in DMEM-high glucose, 20% FBS. When the cells reached 90% confluence, they were treated with differentiation medium (DMEM-high glucose with 10% FBS, 1μg/mL insulin, 1nmol/L T3, 0.125 mmol/L indomethacin, 1μmol/L dexamethasone, 0.5mmol/L isobutylmethylxanthine and 1μmol/L rosiglitazone) for three days, then switched to DMEM plus 10% FBS, 1μg/mL insulin, and 1nmol/L T3 for four days. After 7 days of differentiation, the adipocytes were used for experiments.

For beige adipocyte differentiation, primary preadipocytes were isolated from 6 to 8 weeks old C57BL6/J mice through collagenase digestion. The differentiation was performed by culturing confluent cells in medium (DMEM-high glucose with 10% FBS, 5μg/mL insulin, 1nmol/L T3, 0.125 mmol/L indomethacin, 1μmol/L dexamethasone, 0.5mmol/L isobutylmethylxanthine and 1μmol/L rosiglitazone) for three days, then switched to DMEM plus 10% FBS, 5μg/mL insulin, and 1nmol/L T3 for four days. After 7 days of differentiation, the adipocytes were used for experiments.

To examine the impact of Prg4 on HSL phosphorylation at Ser563, the differentiated adipocytes were incubated with DMEM or DMEM containing Prg4 for 4 hours.

### Western blot

After treatments, cells were washed twice with PBS and extracted with cell lysis buffer (20 mM Tris pH 7.6, 150 mM NaCl, 0.5 mM EDTA, 0.5 mM DTT, 10% glycerol, protease and phosphatase inhibitors). For tissue extracts, the proper amount tissues (i.e. liver etc.) were cut into small pieces and resuspended in SDS-UREA buffer (50mM Tris, 1%SDS, 10% glycerol, 5mM Na_4_P_2_O_7_, 50mM NaF, 5M urea, 5mM EDTA composition). Then, the tissues were homogenized and centrifuged at 13krpm for 10 mins. For total cell lysate and tissue extracts, the protein concentration was determined by BCA kit (Pierce 23225). For cellular fractionation experiments, the cellular fractions were directly dissolved into lysis buffer. Equal amounts of protein were subjected to SDS PAGE electrophoresis and transferred to nitrocellulose membranes (Biorad). After blocking in 5% dry milk, the membranes were incubated with each primary antibody, followed by incubation with a horseradish peroxidase-conjugated secondary antibody. The protein bands were visualized using the ECL detection system (Pierce). The quantification of Western blot was determined with Image J software.

## Protein Palmitoylation Assay

The protein palmitoylation assay has been described (45). Briefly, total cytoplasmic extracts were prepared from cultured cells or mouse tissues. Then the extracts were incubated with blocking buffer (100 mM HEPES, pH 7.6, 2.5% SDS, 1 mM EDTA, 1μL/ml methyl methanethiosulfonate) for 20 min at 42°C, and the proteins were precipitated with 3 volumes of 100% acetone. After centrifugation the protein pellets were dissolved into binding buffer (100mM HEPES, pH 7.6, 1 mM EDTA) plus thiopropyl-Sepharose beads and incubated for 4 h. The beads were washed three times with binding buffer. Then the proteins on the Sepharose beads were eluted with 100 mM DTT and analyzed by western blot with appropriate antibodies.

## Real-time PCR

Total RNA was isolated from mouse tissues or cultured cell by using RNeasy kit (Qiagen), and quantified with a Nanodrop spectrophotometer (ThermoScientific, Wilmington, DE). First Strand cDNA was synthesized with the SuperScript® III First-Strand Synthesis System with random decamers as primers (ThermoFisher). Gene expression was determined by real-time quantitative PCR using a Prism HT7900 instrument with SYBR Green chemistry (Applied Biosystems, Foster City, CA). The primers were indicated in the following: DHHC7F-CTCTGATTCTCTGTGGGCTTC, DHHC7R-GCCTCTCGATCTCTGTTTCATC; DHHC3F-CCACTTCCTGCATTGCTTTG, DHHC3R-TAGCCCATCTTCTCTCTTCCT; 36BP4F-CTGAGTACACCTTCCCACTTAC, 36BP4R-CCGAATCCCATATCCTCATCTG. The procedure was as 94°C for 30s,60°C for 30s, and 72°C for 30s for 40 cycles.

### Proteomics analysis

The liver tissue and plasma were harvested from the high fat diet male mice, then were performed to quantitative proteomics analysis at Shanghai OE Biotech Co., Ltd. (China). The thresholds of fold change (>1.2 for plasma proteomics, >1.5 for liver tissue proteomics) and P-value (<0.05) were used to identify differentially expressed proteins (DEPs).

### Immunoprecipitation assays

Transfected HEK293 cells were extracted with immunoprecipitation buffer (150 mM NaCl, 25 mM Tris, pH 7.6, 0.5 mM EDTA, 10% glycerol, 0.5% NP-40 plus a protease inhibitor mixture). 500 ug of total proteins from the cell lysates were subjected to immunoprecipitation with the corresponding antibodies.

### Statistical Analyses

The quantification of Western blot is carried out with Imagine J. Means ± S.D. were calculated, and statistically significant differences among groups were determined by one-way analysis of variance followed by post hoc comparisons or by two-tailed unpaired Student’s *t*-tests as appropriate, with significance set at *p*< 0.05.

## Acknowledgments

This project is in part supported by The Start Funding of Hebei University (to YS), and the Natural Science Foundation of Hebei Province (YS) (H2021201011), in part supported by the Biological Breeding-National Science and Technology Major Project(XH) (2023ZD0406805).

